# Transcription Factor NF-κB Unravels Nucleosomes

**DOI:** 10.1101/2021.03.18.436050

**Authors:** Tommy Stormberg, Shaun Filliaux, Hannah E. R. Baughman, Elizabeth A. Komives, Yuri L. Lyubchenko

**Affiliations:** Department of Pharmaceutical Sciences, University of Nebraska Medical Center, 986025 Nebraska Medical Center, Omaha, NE 68198-6025; Department of Chemistry and Biochemistry, UC San Diego, 9500 Gilman Dr., La Jolla, CA 92093-0378

## Abstract

NF-κB is a transcription factor responsible for activating hundreds of genes in mammalian organisms. To accomplish its function, NF-κB must interact with DNA occupied by nucleosomes, but how this interaction occurs is unclear. Here we used Atomic Force Microscopy to characterize complexes of NF-κB with nucleosomes assembled on different DNA templates. The assembly of NF-κB-nucleosome complexes leads to a substantial decrease of DNA wrapping efficiency. Mapping of the nucleosomes did not reveal displacement of under-wrapped nucleosomes from their original position, suggesting that unravelling involves dissociation of one or both flanks of the nucleosomes. We discovered two binding modes of NF-κB associated with nucleosome unraveling - NF-κB bound to the nucleosome core and to the DNA flanks and propose models explaining the interaction of NF-κB with the nucleosome. We speculate that NF-κB can function as a pioneer factor enhancing its ability to facilitate rapid transcriptional response to cell stress.

## Introduction

Transcriptions factors (TFs) play an important role in activating gene expression by starting and regulating the copying of DNA to RNA. The nuclear factor-κB (NF-κB) family of TFs regulate the transcription of a diverse variety of genes involved in immune response, inflammation, cell differentiation, and apoptosis(Didonato et al., 2012; Oeckinghaus and Ghosh, 2009; Tripathi and Aggarwal, 2006). NF-κB is a ubiquitous TF, found across mammals and simpler animals, and in nearly all cell types(Ghosh et al., 1998). In mammals, the family is composed of five proteins-p50, p52, p65, RelB, and c-Rel-that form dimers in different combinations to perform different transcriptional functions(Oeckinghaus and Ghosh, 2009). The p65/p50 NF-κB heterodimer (hereafter referred to as NF-κB for simplicity) is the most common complex in the NF-κB family and is especially important in inflammatory response in mammalian cells(Smale, 2011).

NF-κB rapidly responds to inflammatory signals by invading the nucleus from the cytoplasm and binding to DNA target sites represented by the canonical sequence, GGGRNNYYCC, where R is a purine, N is any nucleotide, and Y is a pyrimidine, to start transcription(Brignall et al., 2019; Chen et al., 1998; Pereira and Oakley, 2008). Data has shown, however, that NF-κB is capable of efficiently binding to sequences other than the canonical sequence(Mulero et al., 2019; Wong et al., 2011). However, the mechanism of interaction of with chromatin, and nucleosomes specifically remains unclear. While it is largely agreed upon that nucleosomes can inhibit NF-κB recruitment(Ramirez-Carrozzi et al., 2009; Saccani et al., 2001), conflicting reports have been published on the capability of NF-κB to interact with nucleosomal DNA. An in-depth study utilizing EMSA and footprinting found that the p50 homodimer is incapable of binding a recognition sequence located at the nucleosomal dyad without induced nucleosome destabilization(Lone et al., 2013), while another study demonstrated that the homodimer can indeed invade the nucleosome and bind to an internally held recognition sequence(Angelov et al., 2004). Moreover, it is unclear whether NF-κB is capable of inducing structural nucleosome modulation or eviction.

Here we sought to clarify these questions by using Atomic Force Microscopy (AFM) to analyze NF-κB-nucleosome complexes on the single molecule level with nanometer resolution. We utilized the topographical mapping capability of AFM to image NF-κB on naked DNA and strongly positioned nucleosomes in the presence and absence of NF-κB (p65/p50) heterodimers to determine how nucleosome structure is affected. Our results show that NF-κB unravels the nucleosome structure and we identified two binding modes-interaction with the DNA flanks and a direct interaction between NF-κB and the nucleosome core. We propose that NF-κB binding and nucleosome unraveling suggests it can function as a pioneer factor(Zaret, 2020) that facilitates the transcription processes mediated by transcription factor NF-κB.

## Results

### Interaction of NF-κB with naked DNA substrate

A 377 base pair DNA substrate containing a nucleosome positioning 147 bp Widom 601 sequence motif was used for all experiments (Fig. S1A). The 147 bp Widom 601 sequence was flanked by 113 and 117 bp on either side. The 10 bp NF-κB recognition motif,GGGRNNYYCC, is located near the end of the 601 sequence on the 117 bp flank side. We first used AFM to map NF-κB positioning on the DNA substrate. We prepared the DNA:NF-κB complex at a 1:2 ratio in the protein binding buffer for imaging with AFM. The Widom 601 sequence used contains a NF-κB positioning motif, GGGATTCTCC, located at the end of the 601 Widom sequence (Brignall et al., 2019; Mulero et al., 2019), or ~20 nm from the center of the substrate. Typical AFM images of complexes of NF-κB with DNA are shown in Fig. 1A. Representative zoomed-in images are shown to the right of the large-scale AFM image in Fig. 1B. Positions of the protein appearing as bright features on the DNA filaments are indicated with white arrows. Images in rows (i), (ii), and (iii) depict a single NF-κB, two NF-κB, and three NF-κB dimers in complex with DNA, respectively. The position frequency of NF-κB along the substrate is shown in Fig. 1C. The frequency of position of the first observed protein (closest to the end of the shorter DNA flank) is in blue, while the second and third are in violet and red, respectively. Blue arrows indicate possible positions of the NF-κB binding motif. These data reveal two observations. First, a preference for binding to the end of the DNA is observed. Second, the binding of NF-κB to the DNA is not highly specific to the binding motif-in all three subpopulations, NF-κB is observed to be bound rather randomly across the substrate.

**Figure 1.**
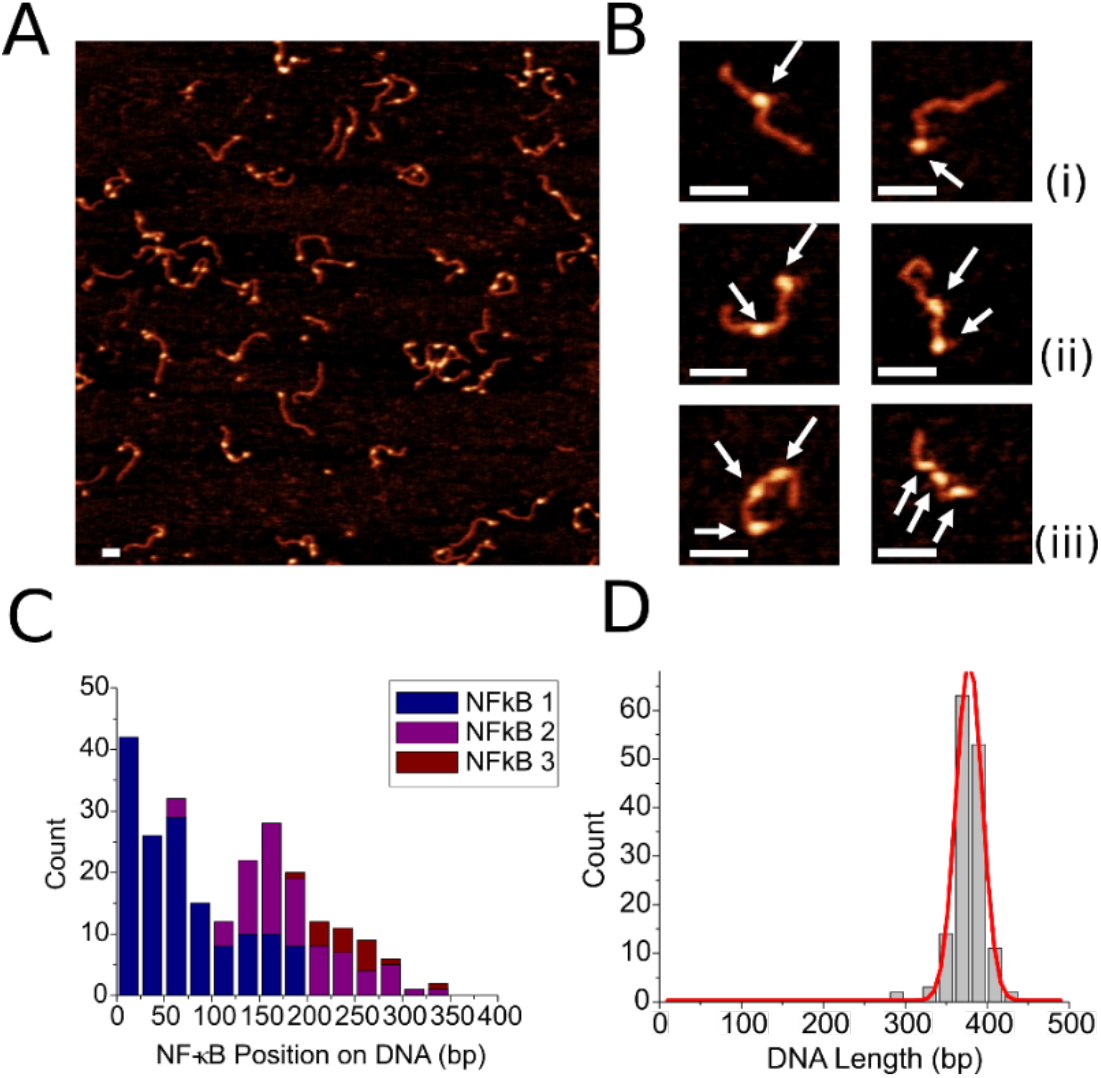
AFM images of NF-κB-DNA complexes. **A.** Representative image, **B.** Zoomed in snapshots of complexes. White arrows point to the protein bound to DNA. (i) DNA with single NF-κB bound, (ii) DNA with two NF-κB bound, (iii) DNA with three NF-κB bound. Bar size is 50 nm. **C.** Frequency of NF-κB position. Blue coloring of bar indicates position of first NF-κB, while purple and red indicate position of second and third NF-κB, respectively **D.** Histogram of NF-κB-DNA complex lengths (*n* = 148) with Gaussian fit indicate a peak centered around 378 bp.

We performed contour length analysis to understand how NF-κB interacts with our substrate. NF-κB-DNA complexes with up to three bound NF-κB dimers were measured from end to end, noting NF-κB position. NF-κB binding to our substrate does not alter the contour length of the DNA (Fig. 1D); thus, NF-κB does not wrap DNA and does not need to be considered when calculating the wrapping efficiency of nucleosomes in complex with NF-κB.

### NF-κB induces unravelling of the nucleosome

Nucleosomes with no NF-κB present in the sample were first imaged and characterized. A representative image of assembled nucleosomes with zoomed in snapshots below is shown in Figure 2A. The bright circular features in the images are the nucleosome cores and the strands spreading from the cores are the unbound DNA of the nucleosome complexes. The snapshots were chosen to represent the different degrees of nucleosome wrapping seen in such samples, with a turn rate of ~1.7 turns being the canonical value. Nucleosome wrapping efficiency defines the length of DNA occupied by the nucleosome core and was calculated by subtracting the measured length of the unbound DNA on each complex from the known length of the DNA substrate, as described in more detail in the Methods section. The histogram of nucleosome wrapping efficiency in these samples, seen in Fig. 2B, demonstrates that the nucleosomes are tightly positioned around the 601 sequence and show a canonical wrapping efficiency of 149 ± 2 bp (SEM). Nucleosomes were also mapped to determine position of the core on the DNA substrate. The results of mapping are shown in Fig. 2C. Each bar represents a single nucleosome complex. The red dots indicate the position the nucleosome begins wrapping from the short end of the DNA substrate. Taken together, these results indicate that the nucleosomes are canonically wrapped and tightly positioned around the Widom 601 sequence, which is consistent with current knowledge and our previous studies on canonically assembled nucleosomes(Stormberg et al., 2019).

**Figure 2.**
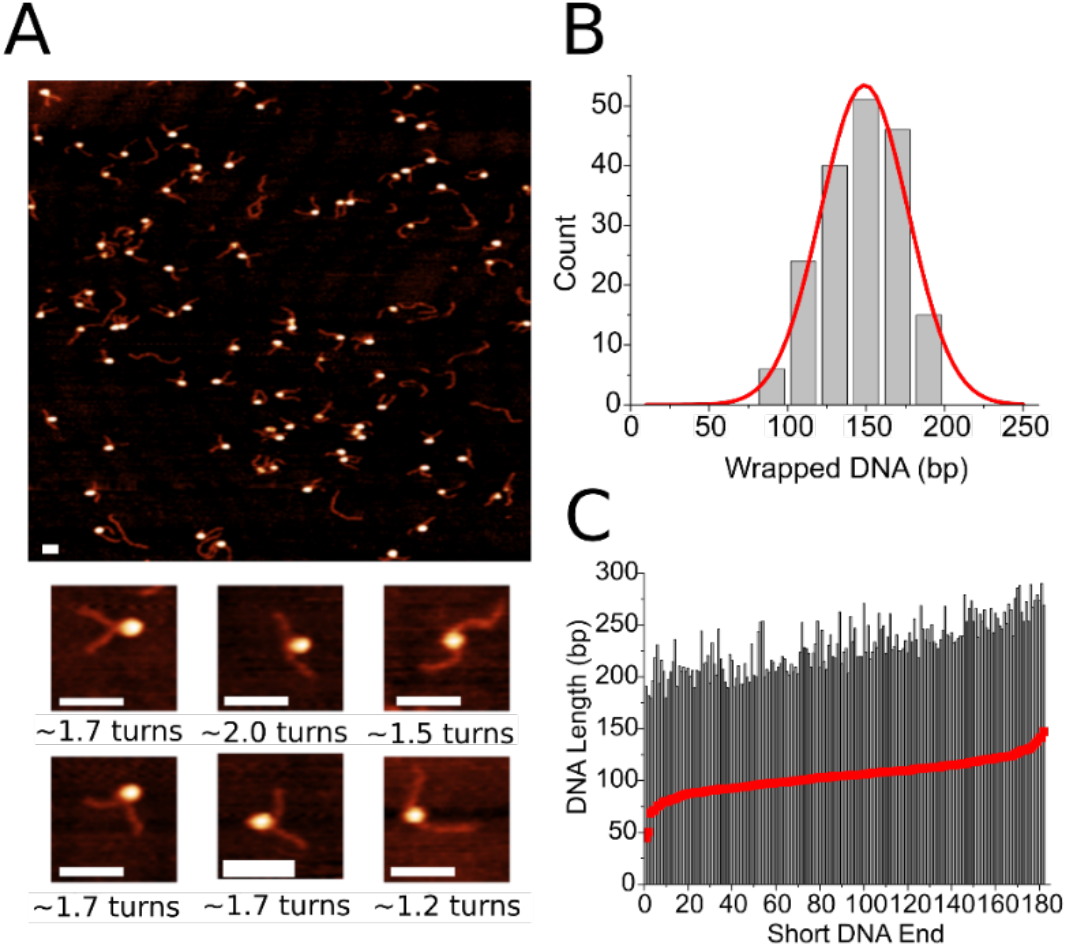
Analysis of nucleosomes assembled on the 601 substrate. **A.** Representative image of assembled nucleosomes with snapshots below. Turns of wrapped DNA are indicated, with ~1.7 turns being the canonical value. Bar size is 50 nm. **B.** Histogram of wrapping efficiency (*n* = 182) reveals a mean wrapping of 149 ± 2 bp (SEM). **C.** Mapping of nucleosome position reveals placement around the 601 positioning sequence. Nucleosomes are indicated by red dots. Each bar represents a single complex.

Next, we studied the effect of the NF-κB protein on the nucleosome structure. NF-κB was incubated for 10 minutes with assembled nucleosomes at a 1:1 ratio before dilution and immediate deposition for AFM imaging. A representative AFM image and selected snapshots can be seen in Fig. 3A. The snapshots show nucleosome-NF-κB complexes in which NF-κB is bound to the DNA flanks (indicated by the blue arrows, nucleosome cores indicated by the red arrows) in different positions along the substrates. NF-κB is observed bound at locations all across the substrate. We calculated the wrapping efficiency of these complexes in the same fashion as above. In determining the length of the unbound DNA for nucleosome wrapping efficiency calculation, DNA bound by NF-κB was considered as free DNA, as no change in the DNA contour length in complexes with NF-κB was detected (Fig. 1C). The histogram of the measured wrapping efficiency in complexes bound by NF-κB on the flanks is shown in Fig. 3B. Nucleosome-NF-κB complexes at a 1:1 ratio demonstrate a decrease in mean wrapping efficiency to 135 ± 3 bp (SEM), corresponding to unravelling of 14 bp compared with the control nucleosome samples. Nucleosomes lacking NF-κB on the flanks, however, demonstrate a mean wrapping efficiency of 146 ± 3 bp (SEM) as shown in Fig. 3C, which is within error of the value of the control.

**Figure 3.**
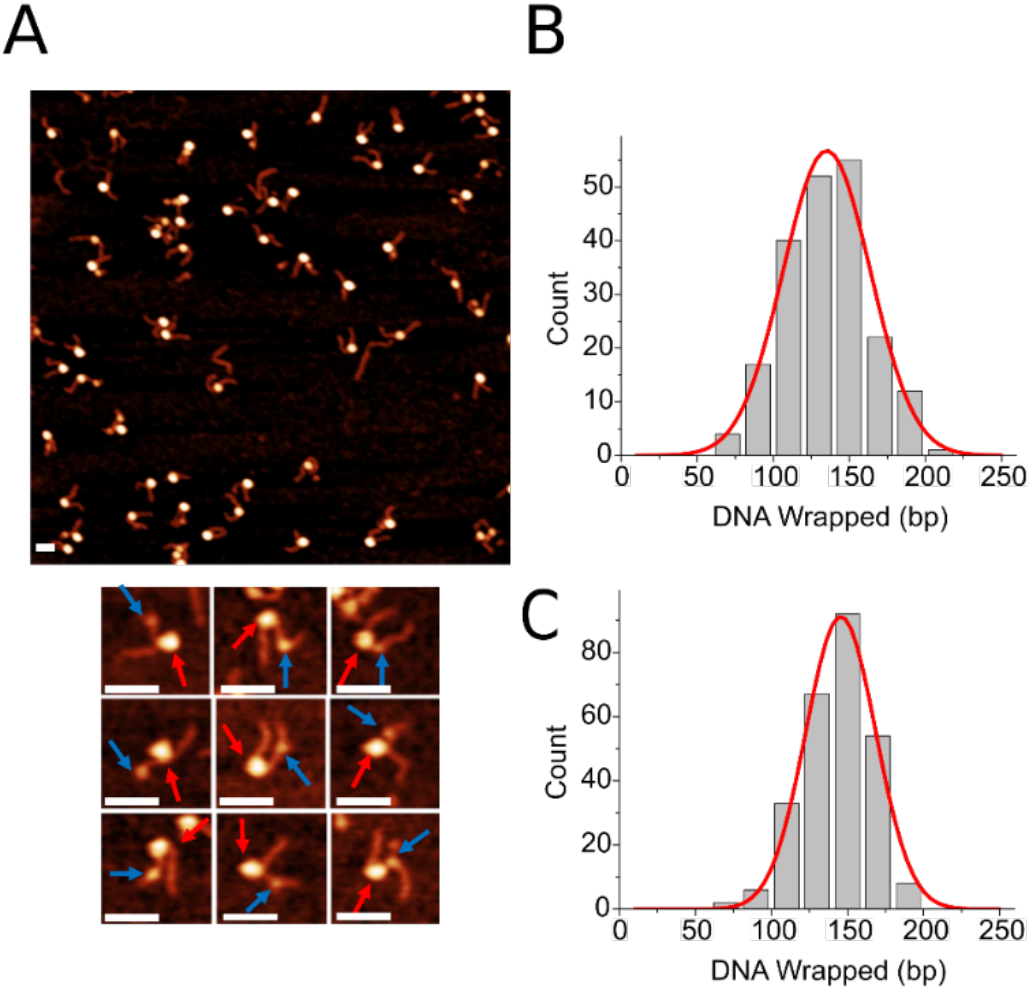
Analysis of nucleosomes incubated with NF-κB at a 1:1 ratio. **A.** Representative image of assembled nucleosomes with snapshots below. Red arrows point to nucleosome core and blue arrows point to NF-κB. NF-κB is shown to position anywhere along the free DNA arms. Bar size is 50 nm. **B.** Histogram of wrapping efficiency (*n* = 203) reveals a mean wrapping of 135 ± 3 bp (SEM) in complexes with NF-κB bound to the flank. **C.** Nucleosomes lacking NF-κB bound to the flanks (*n* = 262) reveal a mean wrapping of 146 ± 3 bp.

To test whether the nucleosome unravelling effect is an inherent property of NF-κB independent of association with the high-affinity nucleosome binding property of the 601 motif, we performed similar experiments with nucleosomes assembled on a DNA template with a nonspecific sequence (Fig. S1B). We in our paper(Stormberg et al., 2019) used plasmid DNA that did not contain a nucleosome binding motif and showed that nucleosomes assembled on this template are morphologically similar to the ones assembled on the DNA containing the 601 motif. The same nonspecific DNA sequence was used here, and the data in Fig. S2A show that wrapping efficiency for nucleosomes assembled on this template is 147 ± 3 bp (SEM). The data obtained for the complexes of these nucleosomes with NF-κB are shown in Fig. S2B. The mean wrapping efficiency of these complexes in the presence of NF-κB was found to be 131 ± 2 bp (SEM). This value is less than the control experiments and even slightly less than the data obtained for 601 nucleosomes. Thus, NF-κB unwrapping is an inherent property of the protein without relation to the DNA sequence.

We performed position mapping of 601 nucleosomes in complex with NF-κB in the same manner as with the control group to determine whether nucleosome unravelling was associated with translocation of the nucleosome away from the 601 motif (Fig. S3). Nucleosomes with NF-κB present on the flanks and those lacking NF-κB on the flanks were both mapped separately. Nucleosome position is indicated in red and NF-κB position in blue-each bar represents a single complex. The mapping indicates that the nucleosome position is not affected by the presence of NF-κB. Nucleosomes are positioned around the region of the Widom 601 sequence as in the control; hence, no translocation is observed.

### NF-κB interaction with nucleosome core

The nucleosome samples incubated with NF-κB contained complexes without the protein bound to flanks, suggesting that complexes of NF-κB associated with the nucleosome core could be there as well. Although the presence of NF-κB bound to the core cannot be identified directly by AFM due to the comparatively smaller size of the protein relative to the nucleosome core, association of NF-κB with the core should increase its apparent size and such complexes can be identified by direct volume measurements of nucleosomes with and without NF-κB.

Nucleosome core volume was determined on a particle-by-particle basis for nucleosomes incubated with NF-κB at 1:1 and 1:2 ratios as well as for nucleosomes with no NF-κB in sample. The resulting histograms are shown in Fig. 4. Nucleosome core volume in samples with no NF-κB have a Gaussian fit centered around 275 nm^3^ (Fig. 4A). For the sample with NF-κB at a 1:1 ratio a bimodal distribution is observed (Fig. 4B). A main peak centered around the same value as the sample with no NF-κB is complemented by a minor peak with a value of 402 nm^3^, with an average fit of 298 nm^3^. This minor peak is converted to a broad shoulder when the NF-κB concentration in experiments is doubled, resulting in a broad Gaussian fit centered around 310 nm^3^ (Fig. 4C). These results suggest that NF-κB is contributing to the volume of the nucleosome core of some complexes.

**Figure 4.**
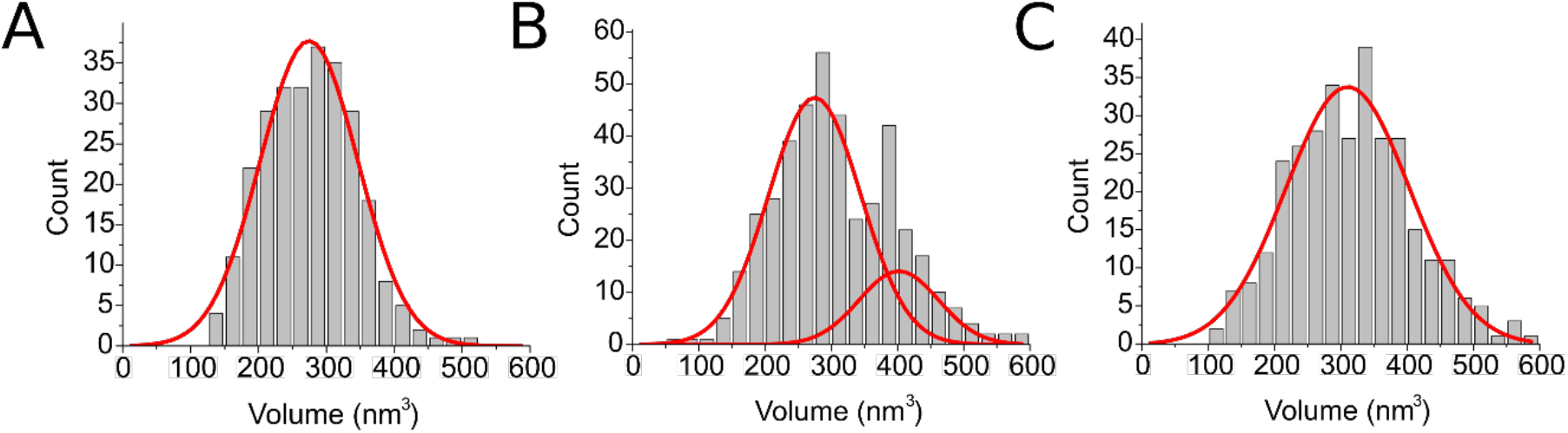
Histograms of nucleosome core volumes with Gaussian fits. **A.** Nucleosomes in samples lacking NF-κB (*n* = 267) have a peak centered around 275 nm^3^. **B.** Nucleosomes incubated with NF-κB at a 1:1 ratio (*n* = 419) reveal a bimodal distribution, with a major peak centered around 275 nm^3^ and a secondary peak centered around 402 nm^3^. **C**. Nucleosomes incubated with NF-κB at a 1:2 ratio (*n* = 314) have a peak volume centered around 310 nm^3^.

Nucleosome core volume is not only dependent on the proteins involved, but also on the amount of DNA interacting with the core. To verify that NF-κB is indeed contributing to the volume and even interacting with the core, we performed a scatter plot comparison and high-volume population comparison of our samples lacking NF-κB against our samples containing NF-κB at a 1:1 ratio (Fig. 5). Graphs in black, 5A(i) and 5B(i), represent the sample population of nucleosomes lacking NF-κB. Graphs in red, 5A(ii) and 5B(ii) represent the sample population of nucleosomes incubated with NF-κB at a 1:1 ratio. Graphs 5A(iii) and 5B(iii) display the merged data of the two sample populations as a single graph. In Fig. 5A, we plotted the core volume of all measured complexes over the wrapping efficiency of the same particles. Each dot represents a single complex. We found that, in samples that contain NF-κB, the core volume of complexes was increased by ~50 nm^3^ on average when wrapping efficiency was considered. This provides further evidence that NF-κB is contributing to the volume of some complexes. In Fig. 5B, we took the subpopulation of complexes with a higher-than-average measured volume (>300 nm^3^) in both samples lacking NF-κB (black) and samples containing NF-κB (red) and plotted a histogram of their respective wrapping efficiencies. Nucleosomes not incubated with NF-κB predictably revealed a mean wrapping efficiency higher than average, as contribution of DNA to the nucleosome core is the primary factor in the volume of the core. However, we found that complexes incubated with NF-κB were consistently under-wrapped when compared with their NF-κB-lacking counterparts. This result indicates that NF-κB directly interacting with the core of the nucleosome particle is also contributing to the partial unravelling of such nucleosomes.

**Figure 5.**
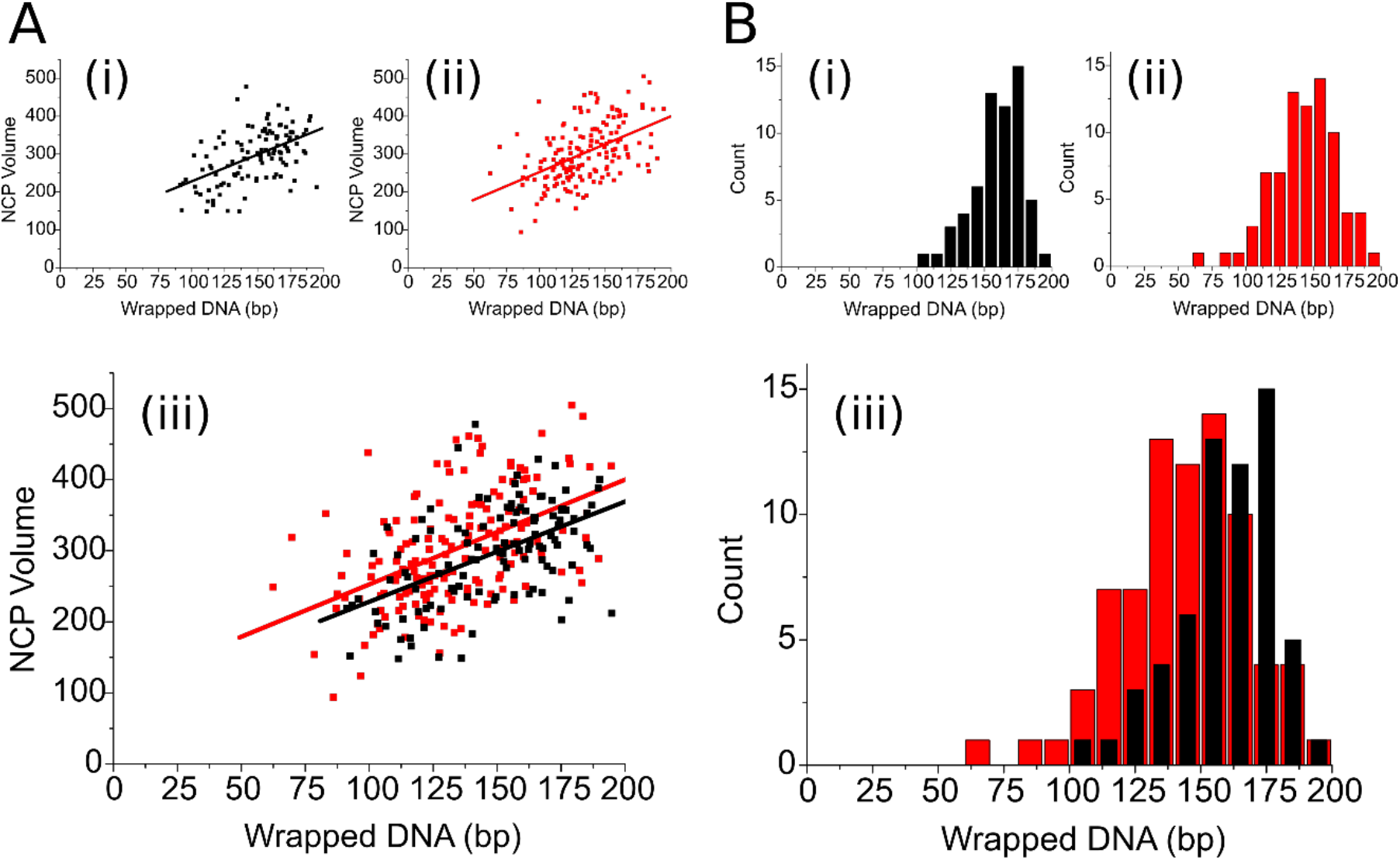
Volume and wrapping comparison of samples lacking and samples with NF-κB present. Graphs in black represent samples lacking NF-κB (A(i), B(i)). Graphs in red represent samples with NF-κB present (A(ii), B(ii)). Merged data of the graphs is presented below (A(iii), B(iii)). **A.** Plotting of nucleosome core particle volume over wrapping efficiency (*n* = 280). **B.** Histograms of the wrapping efficiency of nucleosomes with a nucleosome core volume greater than average (*n* = 139).

We then separated the nucleosome complexes incubated with NF-κB at a 1:1 ratio into two subpopulations-those with NF-κB present on the flank and those without-to distinguish the differences in volume. The results are shown in Fig. S4. The data reveal a higher peak volume for nucleosomes without NF-κB bound to the flanks, but a small subpopulation of oversized nucleosome cores for nucleosomes with NF-κB bound to the flanks is present. These clearly indicate species with one NF-κB bound the flank and another one bound to the core, resulting in the increase of the core volume. Multiple binding of NF-κB to DNA even at the 1:1 ratio is seen in Fig. 1. Together with the wrapping efficiency data (Figs. 3 and 5), we conclude that NF-κB is capable of interacting with the core, and this interaction is associated with the unravelling of the nucleosome.

## Discussion

In this study we sought to characterize the NF-κB-DNA positioning and understand the interaction, or lack thereof, of NF-κB with the nucleosome structure using AFM. We chose the Widom 601 sequence for most experiments which forms a highly stable nucleosome and which contains one consensus binding site for NF-κB. We observed NF-κB bound to the nucleosome and flanking sequences randomly, with little bias towards its consensus binding site consistent with other reports(Wong et al., 2011). The major finding of this paper is that NF-κB decreases the wrapping efficiency of nucleosomes. Importantly, unwrapped nucleosomes were detected for the complexes in which NF-κB was not directly bound to the nucleosome core. We propose two possible models for NF-κB-nucleosome interactions that explains these findings.

### In our first model

interaction of_NF-κB with nucleosome is a dynamic process. After binding the nucleosome core, NF-κB unwraps the nucleosome. NF-κB can dissociate from the core and bind free DNA flanks. However, these species remain unwrapped, suggesting that the interaction of NF-κB with the core changes the structure of the core, so that after dissociation of NF-κB, the core remains unwrapped.

According to our measurements, the mean unwrapping effect of NF-κB is 14 bp for nucleosomes assembled on the Widom 601 sequence and this value is slightly higher for nucleosomes assembled on non-specific M3 DNA template. However, this is a conservative estimate. In fact, each mixture of nucleosomes with NF-κB contain species without bound NF-κB. It is graphically illustrated in Fig. 4B, according to which the population of nucleosomes with bound NF-κB identified by the increased volume is ~20% of nucleosome-NF-κB sample. Given the known wrapping efficiency of nucleosomes without NF-κB, we estimate that the population of species with NF-κB bound to the core can unwrap as much as ~ 30-50 bp. Such a strong unwrapping effect of NF-κB is in line with the data in Fig. 5B(iii). According to this graph, there is a substantial population of species with the nucleosome core bound by NF-κB with wrapping efficiencies below 130 bp. Such an unwrapping effect can be accompanied by a considerable change of the overall structure of nucleosome, perhaps preventing DNA wrapping after the NF-κB dissociation.

### In our second model

NF-κB binds to the nucleosome flank, inducing a downstream loosening of the nucleosome core-bound DNA interaction and allowing NF-κB to interact with the newly unraveled DNA. Our experimental results indicate that NF-κB is able to induce a conformational change in the nucleosome, causing it to adopt a more open conformation. This effect is observed directly when NF-κB is positioned on the free DNA arm of the nucleosome, as illustrated in Fig. 3B. We can see in Fig. 5A that this unraveled state is not always associated with an increased core volume, indicating that NF-κB is not always present at the core at the time of the measurements. Nucleosomes are known to undergo “breathing” in which they transiently wrap and unwrap. The interaction of NF-κB with the DNA flank may shift the equilibrium of this “breathing” toward the unraveled state of the nucleosome by lowering the energy barrier to the more unraveled state. Studies have shown that nucleosomes assembled on 601 are capable of asymmetrically unravelling 10 or more base pairs of DNA under minute levels of tension(Ngo et al., 2015), suggesting that it is reasonable to predict other factors could readily induce similar levels of unravelling, as seen in this study. Interestingly, while preferential binding to the NF-κB positioning motif was not observed on naked DNA, the motif is located within the ~10-20 bp region that more readily unravels on the 601 substrate, suggesting that NF-κB may be positioning there. Binding of NF-κB to the DNA flanks induces a shift to a more open conformation, which in turn allows the protein to more easily bind the nucleosome core. This interaction with the core in turn induces a structural shift which causes the open conformation to become permanent. It is important to note that in this setup the nucleosome is not evicted or translocated, yet it is still destabilized.

Our findings on nucleosomes unraveling by NF-κB is significant in that, coupled with the unravelling effect observed, binding to condensed chromatin is a trait reserved for pioneer transcription factors(Cirillo et al., 2002; Tan and Takada, 2020; Zaret, 2020). While NF-κB is not currently categorized as a pioneer factor, the results in this study suggest that it is capable of acting as one. As suggested above, this direct binding to the core may indeed induce a structural change in the core, permanently inducing a more unraveled conformation of the nucleosome. This finding makes sense in the context that NF-κB is a rapid acting transcription factor responsible for responding quickly to stimuli(Hoffmann et al., 2002). Our results also resolve the questions surrounding chromatin accessibility in the Granulocyte/macrophage colony-stimulating factor (GM-CSF) promoter. Previous studies had shown that nuclear NF-κB levels play a critical role in increased chromatin accessibility at this promoter, but the origin of this effect remained unknown(Holloway et al., 2003). Our results demonstrate clearly that NF-κB by itself can unravel nucleosomes and put to rest the idea that it must rely on other factors for that first step of transcription activation. NF-κB can apparently function as a pioneer factor and by itself unravel the DNA(Zaret, 2020). NF-κB could then recruit the CREB binding protein for subsequent histone modification to stabilize the open chromatin form.

## Materials and Methods

### Preparation of proteins

A pET11a plasmid containing genes for murine p50 (residues 39-350) and p65 (residues 19-321) with an N-terminal 6x-His tag on p50 was transformed into the *E. coli* strain BL-21(DE3) to generate recombinant protein. Cultures were grown at 37°C in 1 L M9 minimal media in the presence of ampicillin to an OD_600_ of 0.6, then transferred to ice for 20 minutes. Expression was induced with 0.2 mM IPTG, and cultures were incubated at 18°C for 16 hours. Cells were harvested, suspended in Lysis Buffer (25 mM Tris, 150 mM NaCl, 10 mM imidazole, 10 mM β-mercaptoethanol at pH 7.5), and 1 mM fresh PMSF and a protease inhibitor cocktail were added. Cells were lysed by sonication, then centrifuged at 12,000 rpm for 45 minutes. The supernatant was loaded onto a 5 mL Ni^2+^-NTA gravity column equilibrated in Loading Buffer (25 mM Tris, 150 mM NaCl, 10 mM imidazole, 10 mM β-mercaptoethanol at pH 7.5) in a 4°C cold room. The column was washed with 100 mL Wash Buffer (25 mM Tris, 150 mM NaCl, 20 mM imidazole, 10 mM β-mercaptoethanol at pH 7.5) and eluted in 30 mL Elution Buffer (25 mM Tris, 150 mM NaCl, 250 mM imidazole, 10 mM β-mercaptoethanol at pH 7.5). The protein was dialyzed overnight against 3 L Dialysis Buffer (25 mM Tris, 150 mM NaCl, 10 mM β-mercaptoethanol, 0.5 mM EDTA at pH 7.5) using SnakeSkin dialysis tubing (7,000 molecular weight cutoff). The sample was loaded onto a MonoS 10/100 column (GE Healthcare) equilibrated in Buffer A (25 mM Tris, 0.5 mM EDTA, 1 mM DTT at pH 7.5), and protein was eluted using a 40 mL gradient from 0-700 mM NaCl. The protein was finally run over a Superdex 200 column in SEC Buffer (25 mM Tris, 150 mM NaCl, 0.5 mM EDTA, 1 mM DTT at pH 7.5).

### Preparation of DNA substrates

The DNA substrates were prepared as we have done previously(Stormberg et al., 2019). Briefly, the DNA 601 substrate used in the experiments was generated using PCR with a pUC57 plasmid vector from BioBasic (Markham, ON, CA). The DNA construct features 147 bp of the strong positioning Widom 601 sequence flanked by plasmid DNA of 113 and 117 bp. The sequence of the construct can be seen in Fig. S5. The random sequence DNA substrate was generated using the same technique as the 601 substrate and is shown in Fig. S6. After the DNA substrates were generated, they were concentrated and purified using gel electrophoresis and separated from the gel using the Gel Extraction Kit from Qiagen (Hilden, DE). DNA concentration was then determined using NanoDrop Spectrophotometer (ND-1000, Thermo Fischer) before being used for nucleosome assembly.

### Nucleosome assembly

Nucleosomes were assembled on both DNA substrates using a gradient dialysis method optimized from our previous research(Stumme-Diers et al., 2018). Recombinant human histone octamers were purchased from The Histone Source (Fort Collins, CO) for use in assembly. Before assembly, histones were dialyzed against initial dialysis buffer (10mM Tris pH 7.5, 2M NaCl, 1mM EDTA, 2mM DTT) at 4C for 1 hour. DNA was then added to the octamer and the initial dialysis buffer was replaced with low salt buffer (10 mM Tris pH 7.5, 2.5 mM NaCl, 1mM EDTA, 2mM DTT) over 24 hours using a dialysis pump. The nucleosomes were then dialyzed for 1 hour against a fresh low salt buffer before being diluted to 300 nM and stored at 4C.

### NF-κB Binding Reaction

NF-κB was diluted to appropriate concentration in NF-κB buffer (25 mM Tris pH 7.5, 150 mM NaCl, 0.5 mM EDTA, 1 mM DTT). Samples of NF-κB bound to free DNA were incubated at a 2:1 protein:DNA ratio with a DNA concentration of 150 nM for 10 minutes at room temperature in sample buffer before deposition. Samples of NF-κB bound to nucleosomes were incubated with a nucleosome concentration of 150 nM at either a 1:1 or 2:1 ratio for 10 minutes before deposition. Nucleosome control experiments were diluted in NF-κB buffer and held at room temperature for 10 minutes before final dilution and deposition.

### Atomic Force Microscopy imaging

Sample preparation for AFM imaging was performed as previously described(Stormberg et al., 2019). Freshly cleaved mica was functionalized with a solution of 1-(3-aminopropyl)-silatrane (APS) for sample deposition(Shlyakhtenko et al., 2012). The samples were diluted from 300 nM to 2 nM in imaging buffer (10 mM HEPES pH 7.5, 4 mM MgCl_2_) immediately before deposition on the functionalized mica. The sample was left to incubate for 2 minutes before being rinsed with water and dried with argon flow. Samples were stored in vacuum and argon before being imaged on Multimode AFM/Nanoscope IIId system using TESPA probes (Bruker Nano Inc, Camarillo, CA). A typical image captured was 1×1 μm in size with 512 pixels/line.

### Data analysis

Data analysis was performed using the methods successfully applied in our lab(Stormberg et al., 2019; Stumme-Diers et al., 2019, 2018). Briefly, DNA contour length analysis was performed by measuring from one end of free DNA to the other using Femtoscan software (Advance Technologies Center, Moscow, Russia). NF-κB position was measured from the end of DNA to the center of NF-κB. Flank measurements for the nucleosomes were obtained by measuring from the DNA end to the center of the nucleosome for both arms. 5nm was subtracted from each measured flank length to account for the length contributed by the histone core. The mean measured value of free DNA on an image was divided by the known length of the given substrate, yielding a conversion unit. All other measurements on each image were divided by the calculated conversion unit to convert measurements in nm to bp. Wrapping efficiency for nucleosomes was calculated by subtracting the combined flank lengths from the known DNA lengths. These methods were used to produce histograms and mapping of nucleosome position and wrapping efficiency. Volume analysis was performed using FemtoScan’s grain analysis feature. Particles to be measured were boxed and automatically outlined. The software then calculated the volume of the outline particles. All graphs were produced using Origin software.

## Acknowledgments

The work was supported by grants to YLL from NSF (MCB 1515346) and NIH (GM096039, GM100156). The authors thank Lyubchenko lab members for useful insights in the data interpretation.

## Supporting Information

**Figure S1.**
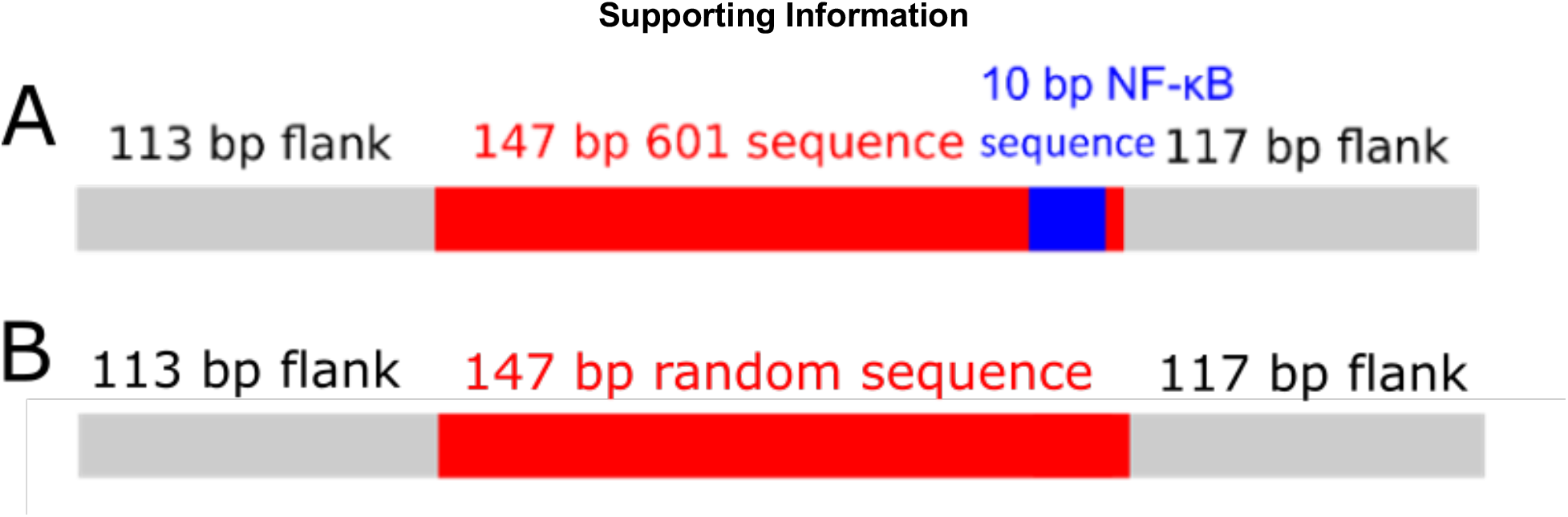
DNA substrates used in experiments. **A.** The substrate is 377 bp long with a centrally positioned Widom 601 motif. The 10 bp NF-κB binding motif is located near the end of the 601 motif on the 117 bp flank side. **B.** The substrate is 377 bp long with a 147 bp random (nonspecific) sequence located in the middle.

**Figure S2.**
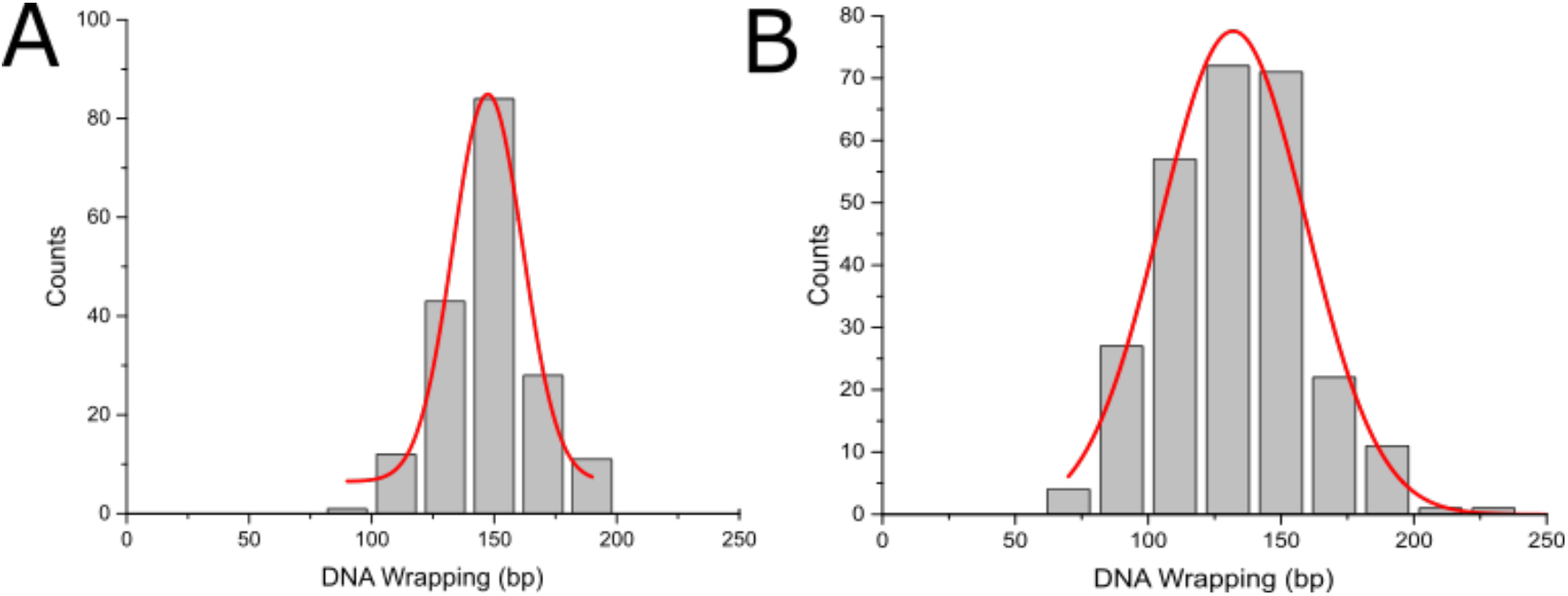
Wrapping efficiency of nucleosomes assembled on the 377 bp nonspecific sequence. **A.** Histogram of wrapping efficiency for nucleosomes assembled on the nonspecific sequence (n = 182) was found to be 147 ± 3 bp (SEM) **B.** Histogram of wrapping efficiency for nucleosomes incubated with NF-κB at a 1:1 ratio (n = 265) reveal a wrapping of 131 ± 2 bp (SEM), indicating an unravelling of 17 bp compared with the control.

**Figure S3.**
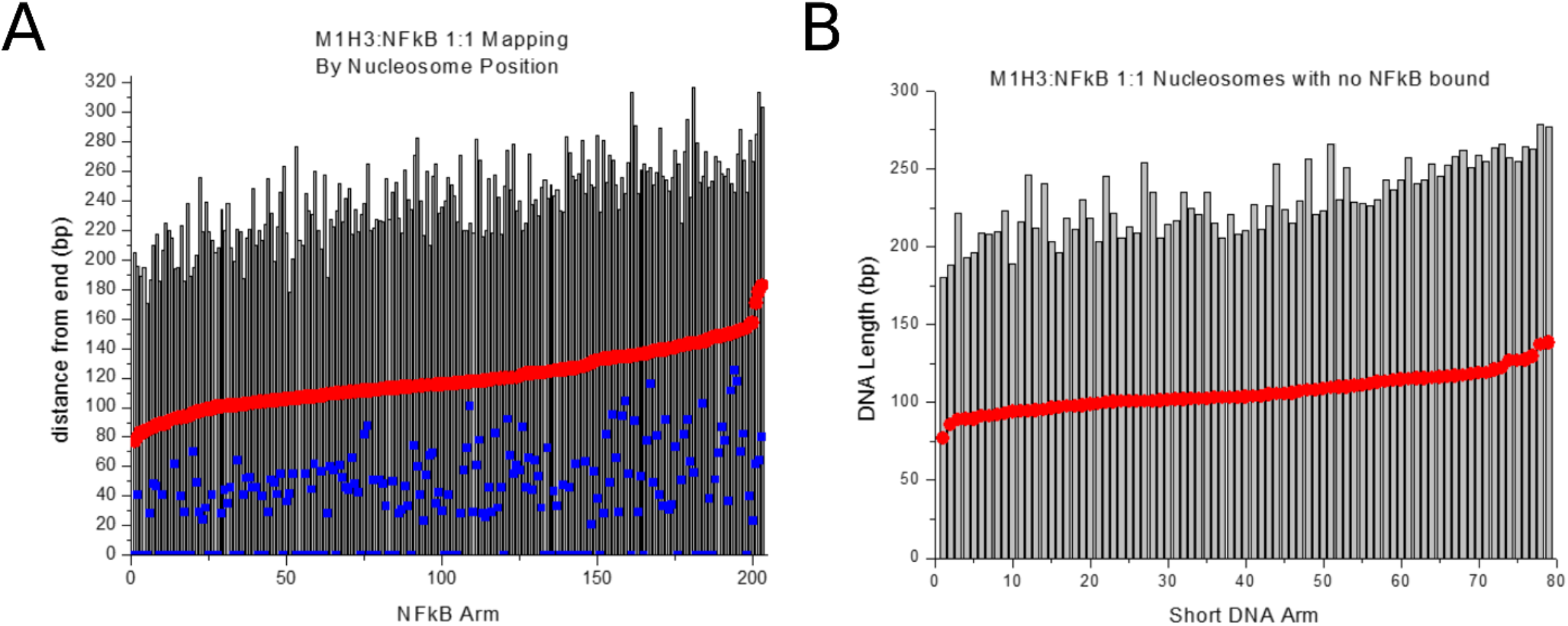
Position mapping of nucleosomes with NF-κB present or absent from flank at 1:1 ratio. **A.** Nucleosomes with NF-κB bound to flanks mapped by position of nucleosome relative to NF-κB. Nucleosomes in red, NF-κB in blue. **B.** Nucleosome mapping with no NF-κB present on flanks.

**Figure S4.**
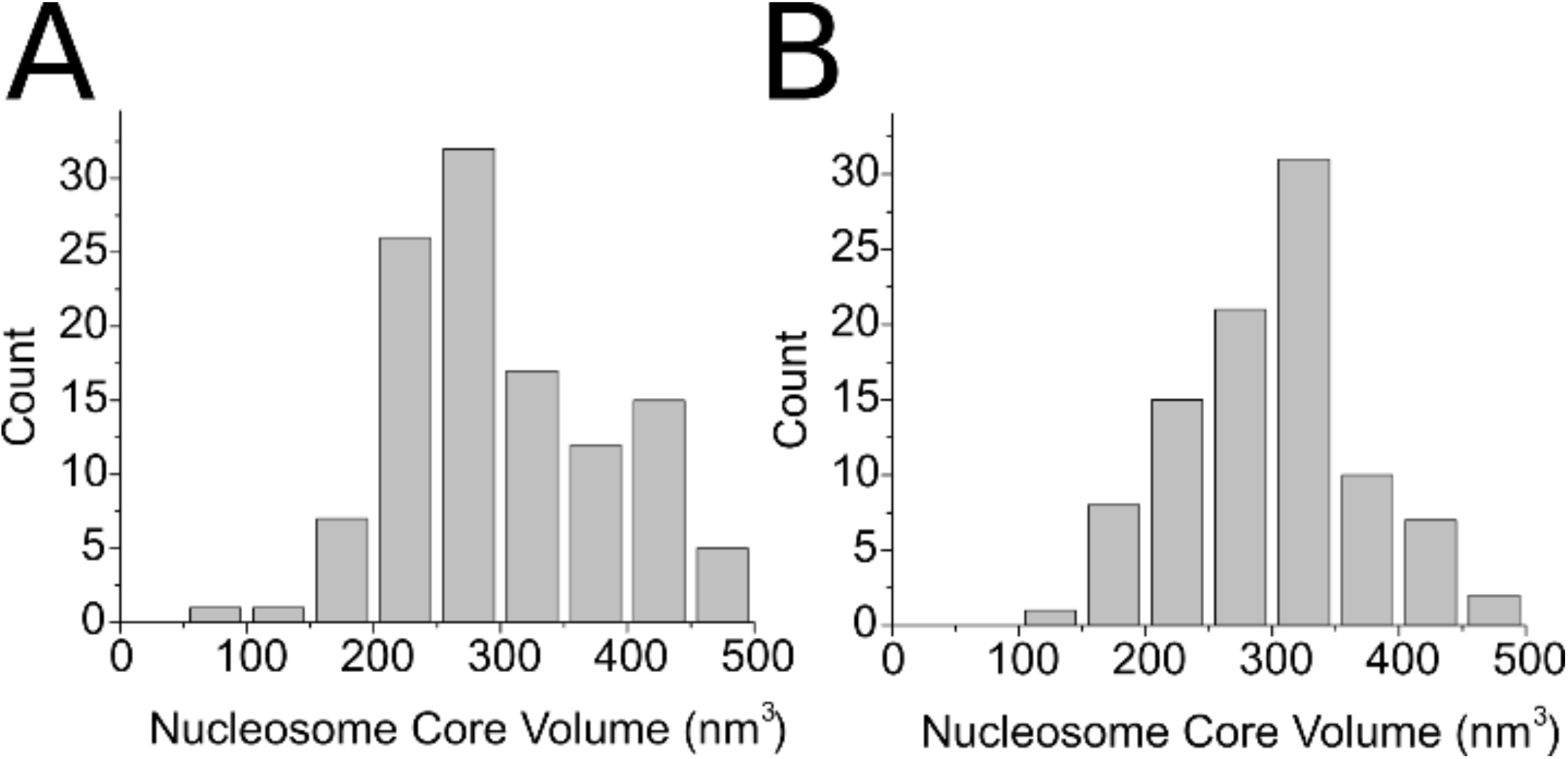
Comparison of nucleosome core volumes of complexes with NF-κB present on a flank **(A)**, (*n* = 95), and absent from the flank **(B)**, (*n* = 116) in samples incubated with NF-κB.

**Figure S5.**
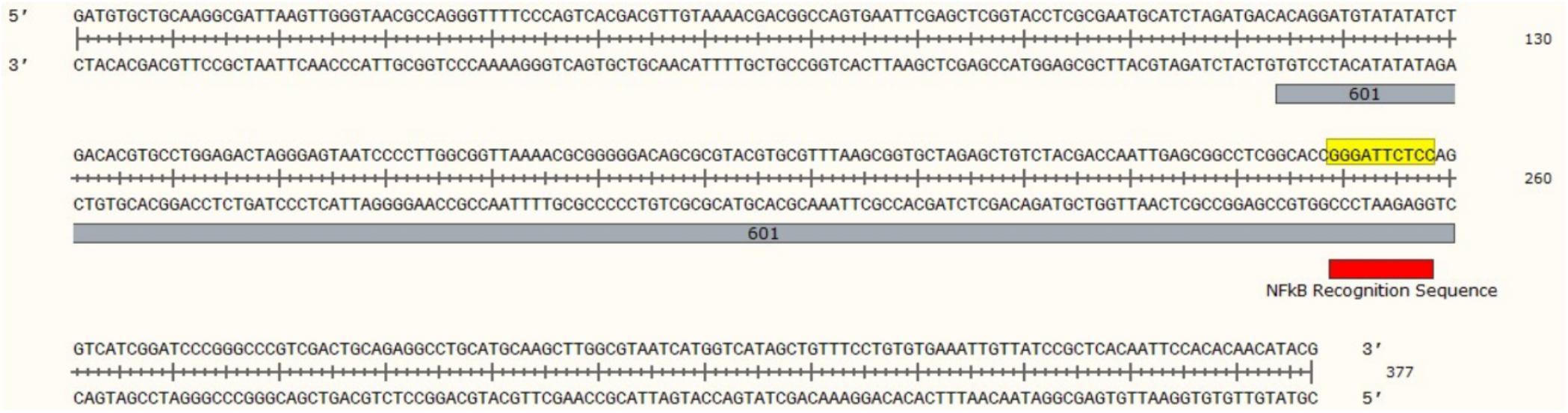
Sequence of 601 DNA construct used for experiments. The DNA construct features 147 bp of the strong positioning Widom 601 sequence flanked by plasmid DNA of 113 and 117 bp and contains the NF-κB binding motif.

**Figure S6.**
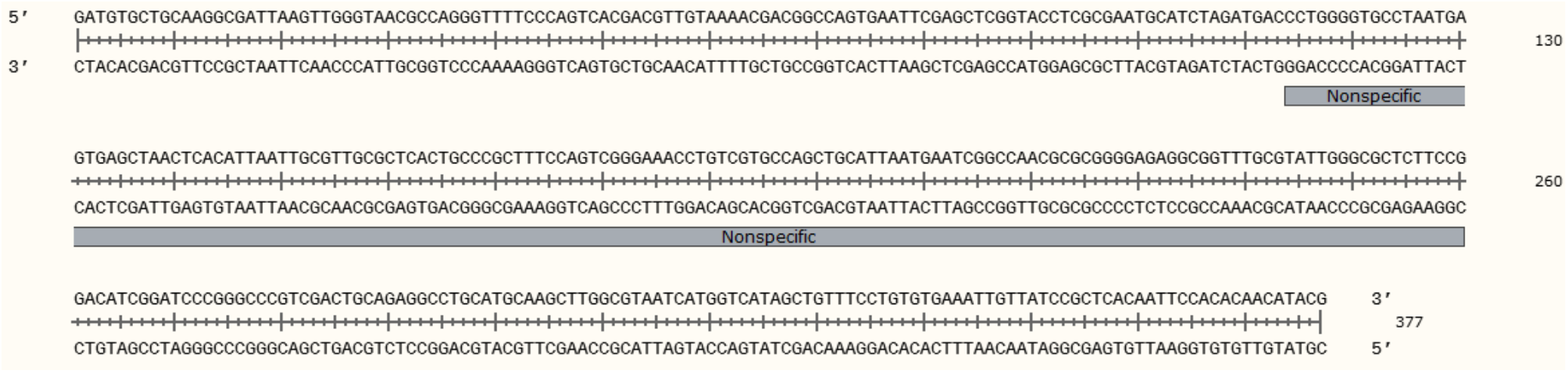
Sequence of nonspecific DNA construct used for experiments. The construct features DNA flanks of identical length and sequence to those used in the 601 construct. In this construct, the 147 bp Widom 601 sequence is replaced by a nonspecific DNA sequence of identical length.

